# Molecular Biology Information Service: An innovative medical library-based bioinformatics support service for biomedical researchers

**DOI:** 10.1101/530071

**Authors:** Ansuman Chattopadhyay, Carrie L. Iwema, Barbara A. Epstein, Adrian V. Lee, Arthur S. Levine

## Abstract

Biomedical researchers are increasingly reliant on obtaining bioinformatics training in order to conduct their research. Here we present a model that academic institutions may follow to provide such training for their researchers, based on the Molecular Biology Information Service (MBIS) of the Health Sciences Library System, University of Pittsburgh. The MBIS runs a four-facet service with the following goals: (1) identify, procure, and implement commercially-licensed bioinformatics software, (2) teach hands-on workshops using bioinformatics tools to solve research questions, (3) provide in-person and email consultations on software/databases, and (4) maintain a web portal providing overall guidance on the access and use of bioinformatics resources and MBIS-created webtools. This paper describes these facets of MBIS activities from 2006-2018, including outcomes from a survey measuring attitudes of University of Pittsburgh researchers about MBIS service and performance.

## Introduction

Recent advancement in molecular technologies such as massively parallel DNA sequencing, microarray platforms, and other high-throughput methodologies, generate substantial amounts of scientific data. Following the successful completion of the Human Genome Project [1,2], initiatives such as the Human Microbiome Project [3,4], the ENCyclopedia of DNA Elements [5], The Cancer Genome Atlas [6], and the 1000 Genome Project [7] continue to generate a massive catalog of biological datasets. In response to this data deluge, bioinformatics software and databases utilizing computational and statistical methods rapidly evolve. Thriving in the current big data-intensive life sciences research environment requires proficiency in bioinformatics tools, which assist with the formulation of new hypotheses, the design of studies to test these hypotheses, and the analysis, interpretation, and validation of experimental results.

The opportunities for experimental biologists to receive bioinformatics training is often limited. Undergraduate and even graduate curricula do not routinely include mandatory bioinformatics classes [8]. Additionally, the bioinformatics resources landscape changes at a rapid pace – the most sought-after resources can quickly become obsolete [9,10]. It is very challenging for biomedical researchers to self-train and stay updated with this moving target.

To help such researchers at the University of Pittsburgh (Pitt), the Health Sciences Library System (HSLS) established the Molecular Biology Information Service (MBIS) in 2002 as an innovative bioinformatics support service. In most biomedical research-intensive institutions, bioinformatics support is typically delivered through departments such as computational biology or biomedical informatics, or facilities such as the sequencing core. Medical libraries traditionally support biomedical research by providing access to journals and books, procuring licenses for electronic resources, providing instructional workshops, and efficiently delivering digital content. As shared-use facilities, libraries can leverage their operational infrastructure to provide a bioinformatics-focused information service by incorporating molecular databases and software into their collections and offer training on the use of these resources.

HSLS was an early pioneer in the implementation of a health sciences library-based bioinformatics support service. The first library to offer a bioinformatics service was the University of Washington Health Sciences Library (UW). The UW service was initiated in 1995 and directed by a PhD scientist [11]. HSLS emulated the UW program in 2002 by hiring a PhD molecular biologist and developing a four-facet service with the following goals: (1) identify, procure, and implement commercially-licensed bioinformatics software, (2) teach hands-on workshops using bioinformatics tools to solve research questions, (3) provide in-person and email consultations on software/databases, and (4) maintain a web portal providing overall guidance on the access and use of bioinformatics resources and MBIS-created webtools.

MBIS has been well-received by the Pitt research community and is now in its second decade of service. Program activities recorded from implementation to 2006 were previously described [12,13]. Several other university libraries have also embraced this type of bioinformatics-focused specialized service as a means to connect with their own biomedical research communities [14–16]; a 2006 special issue of the Journal of Medical Library Association describes a few of these programs in detail [17–19].

MBIS workshops initially focused on information retrieval and searching strategies for molecular databases, and the original HSLS-licensed software products were intended for low-throughput DNA and protein sequence analysis. However, the attention of the research community has shifted to massively parallel sequencing technologies, aka Next Generation Sequencing (NGS). The advances in NGS technology make it is easier, cheaper, and faster to generate huge volumes of sequencing data. Bioinformatics software is routinely developed to analyze these complex, large datasets: RNA-Seq for gene expression studies [20], Exome-Seq for variant detection [21], and ChIP-Seq [22]and ATAC-Seq [23] for epigenomic experiments. Application of these computational tools is critical for experimental scientists to uncover molecular mechanisms underpinning intricate biological processes and diseases. Proficiency with NGS software requires appropriate training with sufficient access to leading tools [24]. As Pitt researchers increasingly began to request assistance with NGS data analysis, MBIS refined its approach by developing collaborative partnerships with other university units and expanding services with the addition of new resources and personnel.

This paper describes the four facets of MBIS activities from 2006-2018, as well as outcomes from a survey measuring attitudes of Pitt researchers about MBIS service and performance. Our intention is that universities with libraries considering or currently providing a specialized bioinformatics service might glean valuable information from the approaches taken and lessons learned by the Pitt HSLS MBIS program.

### University & Library Environment

HSLS thrives in a rich environment of biomedical education, research, and clinical practice at Pitt. As of the 2018 fall term, total undergraduate/graduate student enrollment was more than 28,000, with over 5,000 full or part-time faculty members, and over 700 postdoctoral or research associates [25]. In terms of federal funding, Pitt ranks fifth in competitive grants awarded by the U.S. National Institutes of Health and ninth in federal science and engineering funding, according to a report by the National Science Foundation [26].

Positioned under the Pitt Office of the Senior Vice Chancellor for the Health Sciences, HSLS primarily supports the university’s six schools of health sciences: dental medicine, health and rehabilitation, medicine, nursing, pharmacy, and public health. HSLS staff currently includes 29.2 faculty librarians and 26 paraprofessional and technical staff. The MBIS program started in 2002 with one faculty librarian, added a second faculty librarian in 2007, and a third in 2018.

### MBIS Survey

To evaluate the effectiveness of the MBIS program, an online Qualtrics survey was administered during a six-week period in early 2018. The survey was advertised via numerous methods: MBIS blog post [27], HSLS website post, MBIS listserv notification, direct email invitations, and during MBIS workshops. Participation in the survey was voluntary and anonymous. The survey questionnaire and results are available on figshare [28].

The survey consisted of 33 questions organized in six categories: Demographics, Software, Instruction, Website, Service, and Outreach. A total of 135 participants started the survey, but 13 respondants completed only the demographic questions and were subsequently eliminated, leaving 122 responses for analysis. The majority of responses were from the School of Medicine (66%, n = 81), followed by the Graduate School of Public Health (16%, n = 20). Almost half of the respondents were faculty members (47%, n = 57), followed by students (25%, n = 30), staff (16%, n = 19), and postdocs (11%, n = 13). Among the survey respondents, 50% (n = 80) use Windows operating system, 38% (n = 61) use Mac, and 12% (n = 19) use Linux; respondants could select more than one operating system.

### MBIS Bioinformatics Software

MBIS identifies and procures network licenses for commercially-available bioinformatics software, implements license servers, and facilitates access free-of-charge to the Pitt research community. The library’s operating budget supports most licensing costs, with supplementary funding provided by Pitt’s Institute for Precision Medicine[29]. Available software programs are listed on the MBIS website [30]; upon registration, users receive an email with detailed access instructions. HSLS licenses 17 proprietary software packages as of January 2019; the software list, vendors, and year of initial licensing is shown in Table 1.

**Table 1.** HSLS MBIS-licensed software

| **Software/Database** | **Vendor** | **Software Analysis Classification** | **Initial Licensing Year** | **Number of Registrations Jan 2019** | **Publications by Pitt Researchers Citing the Software (Google Scholar\*)** |
| --- | --- | --- | --- | --- | --- |
| BaseSpace Correlation Engine (NextBio) | Illumina | Gene Expression | 2012 | 515 | 26 |
| Biomedical Genomics Workbench | QIAGEN | Next Gen Seq | 2016 | 359 | 2 |
| CLC Genomics Workbench | QIAGEN | Next Gen Seq | 2010 | 819 | 96 |
| CLC Main Workbench | QIAGEN | Seq Manipulation | 2010 | 620 | 11 |
| ePath3D | Protein Lounge | Scientific Illustrations | 2015 | 246 | 2 |
| GeneSpring GX | Agilent | Gene Expression | 2007 | 244 | 157 |
| GenomeTrax** | BIOBASE | Mutations | 2010 | N/A | 0 |
| Human Gene Mutation Database | QIAGEN | Database | 2012 | 211 | 74 |
| Ingenuity Pathway Analysis | QIAGEN | Pathways | 2006 | 883 | 595 |
| Ingenuity Variant Analysis | QIAGEN | Mutations | 2016 | 151 | 2 |
| Lasergene | DNASTAR | Seq Manipulation | 2004 | 319 | 178 |
| MetaCore | Clarivate Analytics | Pathways | 2007 | 273 | 85 |
| Partek Genomics Suite | Partek | Gene Expression | 2009 | 315 | 109 |
| Protein Center*** | Proxeon | Mass Spec | 2008 | N/A | 9 |
| Sequencher | Gene Codes | Seq Manipulation | 2004 | 325 | 241 |
| SnapGene | GSL Biotech | Seq Manipulation | 2017 | 163 | N/A |
| Transfac/Match and Proteome | GenXplain GmbH | Gene Regulation | 2002 | 344 | 182 |
| Vector NTI Advance | ThermoFisher Scientific | Seq Manipulation | 2002 | 425 | 113 |

\* Publications with at least one University of Pittsburgh-affiliated author; search time frame = 2018 through the Initial Licensing Year + 1
\*\* Discontinued in 2015
\*\*\* Discontinued in 2011

Prior to 2006, HSLS-licensed resources mainly included DNA/protein sequence manipulation software such as *VectorNTI* [31], *Sequencher* [32], *Lasergene* [33], as well as *Ingenuity Pathway Analysis* [34]. In response to user demand, MBIS offerings shifted over the past decade from information retrieval and small-scale sequence analysis tools to sophisticated bioinformatics software capable of handling large NGS datasets.

Many freely-available open source NGS analysis tools use a Command-Line Interface and require some coding experience to operate. Unfortunately, experimental biologists often lack such experience, which creates a significant barrier for tool usage. In response, several vendors offer commercial Graphical User Interface (GUI) software packages that are biologist-friendly and easy to operate [35]. However, GUI-based software is often prohibitively expensive for individual use and typically requires high-performance computers to run the analyses.

To assist Pitt researchers with these issues, HSLS began licensing NGS analysis software and developed strategic partnerships with the Pitt Center for Research Computing (CRC) [36] for advanced computing services, as well as with the Institute of Precision Medicine [29] for funding support in 2011. HSLS negotiated an institutional license with unlimited concurrent users for *CLC Genomics Workbench* [37] along with a license for *CLCbio Genomics* server software. *CLC Genomics Workbench* is a software suite for NGS data analysis such as RNA-Seq, ChIP-seq, Whole Genome Sequencing, Exome Sequencing, and Metagenomics. It is a downloadable GUI software that runs on Windows, Mac, and Linux operating systems. The *CLCbio Genomics* server software is installed at the CRC High Performance Computing cluster (HPC). Together these tools allow users to connect their workstation (desktop or laptop) to a multi-core HPC and transparently migrate data to and run analyses on the cluster.

Software usage is tracked by (1) counting the number of registrants, (2) analyzing software usage log files, and (3) identifying peer-reviewed published articles that cite specific software used, typically in Methods sections. To measure the impact of our software resources on scientific research output, we searched Google Scholar to identify published and preprint articles by Pitt researchers that cite the use of HSLS-licensed software tools. Examination of software registration and citation data reveals frequent usage by the Pitt research community (Table 1).

MBIS survey results indicate overall user satisfaction with software-related services. Seventy-five percent of respondents are very satisfied (n = 91) with the range of software programs offered, 19% indicate that some of what they need is offered (n = 23), and no one expressed dissatisfaction with current offerings. When asked how satisfied they are with the software registration process, including delivery of access instructions, 64% think the process works smoothly (n = 78), 23% indicate that the registration process is fine, but the access instructions could be improved (n = 28), and 2% express frustration (n = 2). Regarding access issues, 48% report no access problems (n= 58), 31% sometimes have access problems (n = 38), and only 2% often experience access problems (n = 3). When asked whether they reference any MBIS commercially-licensed software tools in their publications, 14% indicate yes (n = 17), 46% report “not yet but will soon” (n = 56), and 40% indicate no (n = 49).

The university-wide licensing of proprietary bioinformatics software by HSLS has proven to be very beneficial. It results in significant cost savings by relieving the financial burden of individual labs to license and manage essential commercial bioinformatics software packages [35]. Software is quite expensive for individual research groups, and vendors routinely release software updates at an additional expense. Bioinformatics software usage by individual research groups is normally a project-based process. In our experience, CLC Genomics Workbench is extensively utilized during transcriptomics data analysis, but then rarely used until a similar project starts in the lab. Since HSLS covers yearly maintenance for all licensed software, users can always access the latest version of the software, free of cost.

### MBIS Bioinformatics Training

MBIS provides bioinformatics instruction by teaching guest lectures in graduate and undergraduate for-credit classes, as requested by individual departments or labs, and through in-library workshops. Hands-on workshops covering contemporary bioinformatics topics are typically held in a library classroom equipped with 24 desktop computers. There are three types of library-based workshops: (1) 2 to 3-hour workshops taught by MBIS faculty librarians, (2) workshops supported by software vendors, during which field application scientists showcase salient features of HSLS-licensed tools, and (3) classes taught by invited experts, such as software developers or expert users either from the Pitt community (e.g., Department of Biomedical Informatics, Genomics and Proteomics core facilities) or outside institutions (e.g., University of California-Santa Cruz Genome Browser team, the Galaxy project from Pennsylvania State University).

Workshops are announced via numerous methods: MBIS blog posts [27], MBIS listserv notifications, HSLS newsletter postings, library and health sciences web calendars, monitor slides, and posted flyers. Each workshop has an accompanying online information portal with PowerPoint slides, relevant references, and links to dataset downloads, online tutorials, and additional software/databases [38].

MBIS staff select workshop topics with the aim of helping experimental scientists gain the necessary competencies to be able to use bioinformatics tools to solve their research questions. Most workshops follow a similar format: (1) start with a published dataset imported from a public repository, then (2) build a workflow by combining commercially-licensed and open access software to analyze the dataset. This teaches best practices for data analysis, including recommended parameter settings and interpretation of results. Attendee feedback is generally collected after each workshop and used to modify workshop content or flow as needed.

Hands-on bioinformatics workshops are well received and are an effective means of increasing MBIS program visibility within the Pitt research community. Figure 1 lists MBIS workshops offered during 2007 through 2018 and Figure 2 provides annual workshop statistics: total attendance for all workshops, number of times offered, average number of attendees per session, and number of unique topic workshops. Prior to 2012, popular workshop topics included: Introduction to National Center for Biotechnology Information (NCBI) and European Bioinformatics Institute (EBI) databases for gene/protein-centric information retrieval, DNA/protein sequence analysis using open-access tools and the commercially-licensed *VectorNTI* software suite, sequence similarity searching using *NCBI-BLAST*, biological pathway analysis with *Ingenuity Pathway Analysis* and *MetaCore*, genome sequence browsing, and microarray data analysis.

**Figure 1.**
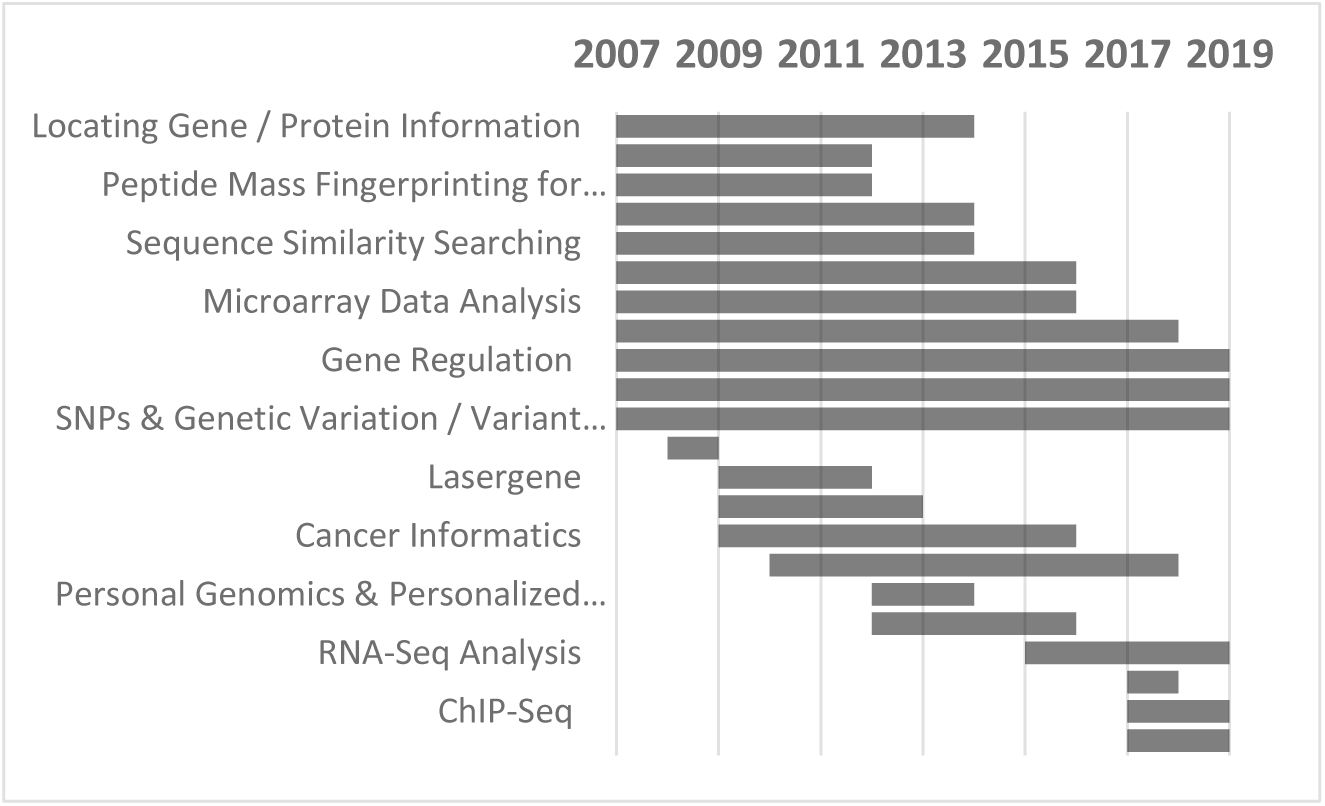
HSLS MBIS hands-on bioinformatics workshops from 2007 through 2018. Bar length indicates the time span of when a workshop was first through last offered.

**Figure 2.**
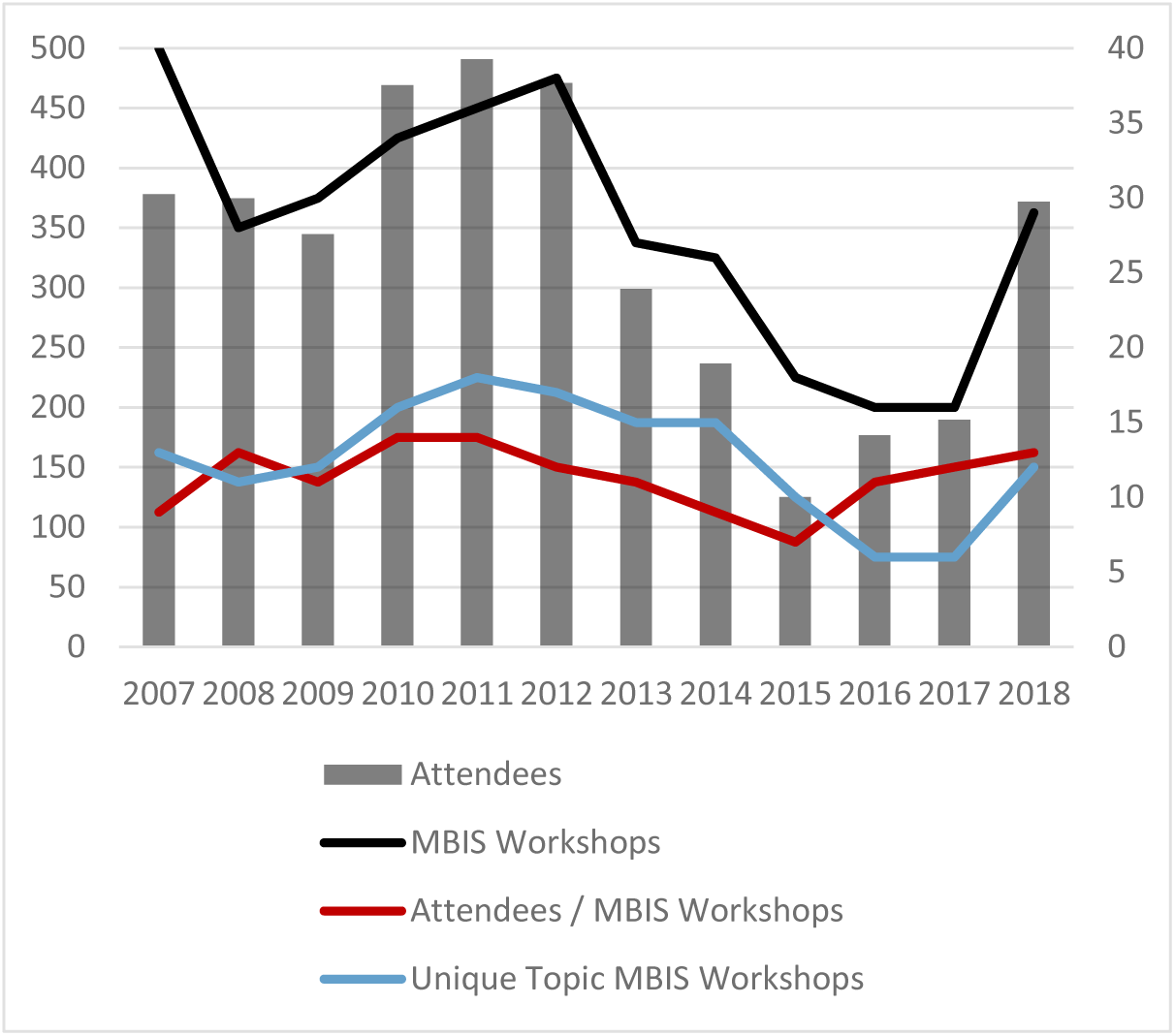
HSLS MBIS hands-on bioinformatics workshops from 2007 through 2018. Thick bars correspond to Left Y axis; thin lines correspond to Right Y axis. Grey = total number of workshop attendees each year; black = number of all workshops offered each year; red = number of attendees divided by number of all workshops offered each year; blue = number of unique topic workshops offered each year.

Starting from 2012, we observed a trending decrease in attendance for workshops covering gene/protein information retrieval, BLAST searching, and microarray data analysis, with a significant drop in the average number of attendees per class in 2015 (Figure 2). We hypothesize that the growing prevalence of online tutorials and videos addressing those topics may have contributed to this drop, as such methods supply researchers with alternative means of training. Alternatively, many of these approaches had by then become standard research practices, so researchers may have moved on to requiring training in different methodologies.

By tracking patron requests along with the development of novel research technologies, we observed an increasing demand for training on Next Generation Sequencing (NGS) analysis, particularly high-throughput bulk RNA-Seq data analysis and variant detection/analysis from data generated through Whole Genome Sequence (WGS) or Whole Exome Sequencing (WES) projects. In 2013, with the availability of a combined software-hardware solution for NGS analysis through the licensing of *CLC Genomics Workbench* along with a strategic partnership with the CRC, MBIS was well-positioned to offer classes on advanced bioinformatics topics dealing with RNA-Seq and variant analyses. NGS workshops on the use of *CLC Genomics Workbench* began in 2014, with the addition of RNA-Seq analysis in 2015, followed by variant analysis in 2016, and ChIP-Seq analysis in 2017. Of note, workshop attendance over 2016-2017 rose after we reevaluated and updated our class topics.

The NGS workshops have been immensely popular from the start, resulting in full classes. To meet the strong demand, in March 2017 MBIS began offering RNA-Seq workshops once a month. Prior to teaching NGS workshops, MBIS workshops were offered on a first-come-first-served basis; the additional demand also prompted the introduction of a class registration process. Interest in these workshops is consistent, and waitlists are not uncommon.

In bioinformatics, the recommended practice is to analyze the same dataset using multiple software packages and then compare the results [39]. As the use of NGS becomes increasingly common for biomedical research, MBIS continually receives requests for training on open source NGS tools. We responded to this unmet need by capitalizing on our experience with the CRC in the implementation of *CLC Genomics Workbench* by also installing *Galaxy*, an open-source web-based GUI-enabled platform for NGS analysis. In 2017, MBIS began teaching RNA-Seq analysis workflow using open-source software with *Galaxy*, and in 2018 added transcription factor and histone ChIP-Seq analysis using *Galaxy* tools.

The addition of these NGS workshops obliged MBIS to discontinue workshops with lower attendance in order to cope with the limited time and classroom space available each semester. As a result, workshops on topics such as DNA/protein information retrieval, sequencing similarity searching, and DNA/protein sequence analysis tools are no longer listed in the workshop roster (Figure 1). According to the MBIS survey, researchers do want specialized workshops covering statistics and data science. MBIS and the HSLS Data Services team [40] therefore developed a partnership with a faculty expert from the CRC to teach Python workshops in 2017, and a workshop on using R for genomics begins in 2019.

MBIS survey results indicate overall satisfaction with education-related services. 68% (n = 81) report that the level of training in workshops is just right, 7% (n = 8) think that too much information is covered, and 5% (n = 6) think the opposite. Regarding the application of information learned in an MBIS workshop (multiple responses allowed), 31% (n = 58) indicate it improved data capture and/or analysis, 14% (n = 27) suggest it changed some aspect of their workflow, 12% (n = 22), 11% (n = 20), and 10% (n = 18), respectively, apply knowledge gained to publications, presentations, and grant applications. Regarding the relevance of workshop topics to their needs, 80% find them very relevant (30%, n = 36) or relevant (50%, n = 60), with only 3% (n = 3) indicating that topics are irrelevant. For a question asking survey respondants to rank the order of their preferred mode for workshops, strongly in first place is in-person/hands-on at 70% (n = 75), followed by tutorials (17%, n = 17) or videos (14%, n = 13), with live webinars in a distant last place (6%, n = 5). For workshop duration, two hours is slightly more favored 59% (n = 53) over the typically offered three hours timespan (41%, n = 37). It can be a challenge determining an appropriate day/time for workshops, but the survey results indicate that the current day and start time for workshops is modestly preferred: Wednesdays (54%, n = 45) at 1 pm (60%, n = 68).

### MBIS Bioinformatics Consultations

Biomedical researchers regularly contact MBIS for assistance. These interactions fall into two general categories: (1) prompt responses to relatively simple email questions, often on licensed software access issues and (2) 1-2 hour in-person consultations providing individualized training on complex bioinformatics software applications; multiple sessions are often necessary to fully resolve these queries. Figure 3 lists the annual number of consultations from 2010-2018 for each of these interaction types. The increased number of in-person consultations beginning in 2016 is associated with the advent of NGS data analysis software and workshops.

**Figure 3.**
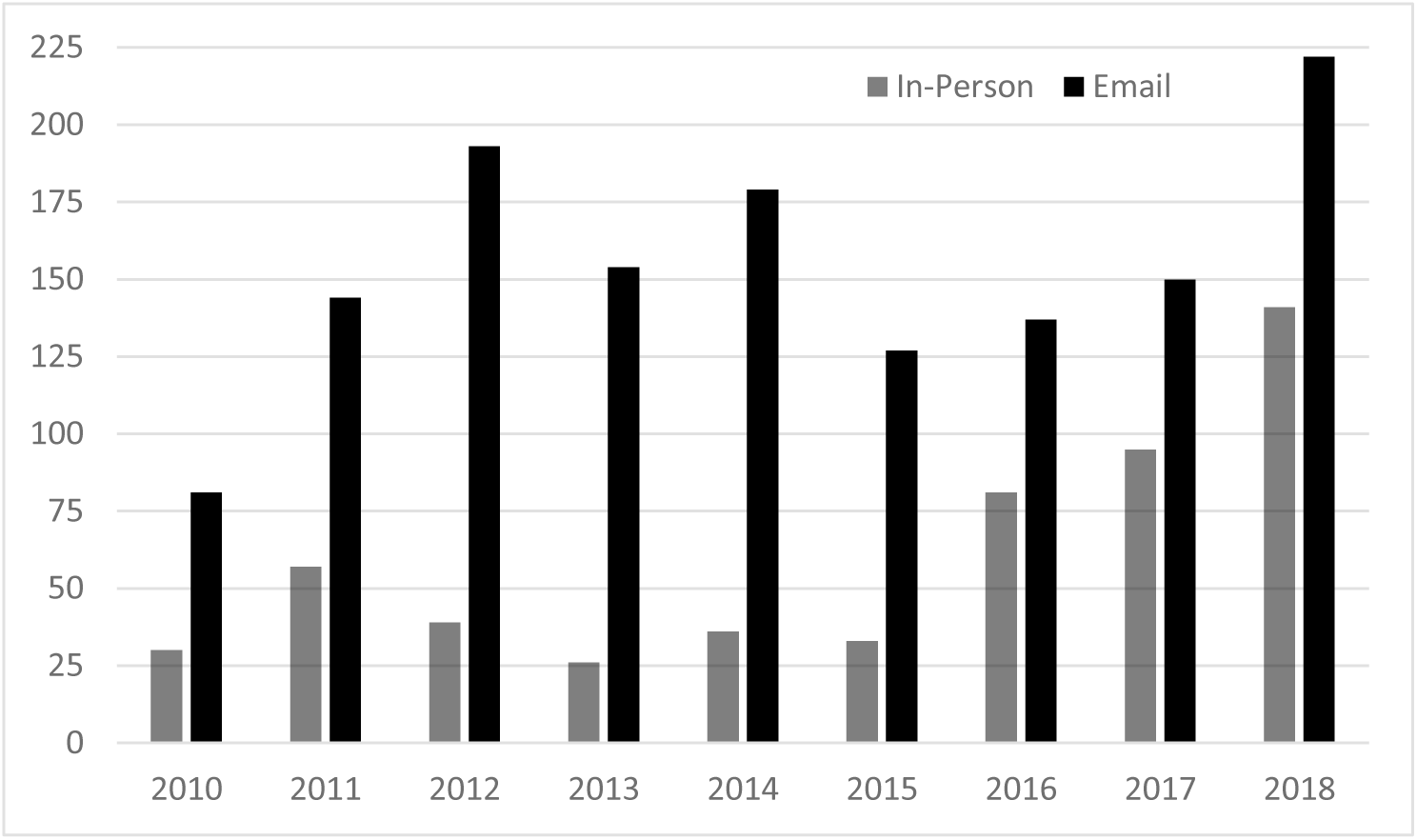
HSLS MBIS consultations from 2010 through 2018. Grey = number of in-person 1-2 hour consultation sessions; black = number of email interactions, including software troubleshooting.

The analytical workflows for NGS software are complex for many researchers, and rigor and reproducibility in data analysis is a growing concern. MBIS workshops provide a solid overview of the multi-step process, but additional training is often necessary to ensure competence. In response, MBIS collaborated with the Genomics Analysis Core of the Department of Biomedical Informatics (DBMI-GAC) to hire a GAC analyst to provide comprehensive assistance with *CLC Genomics Workbench* data analysis. The analyst started on a part-time basis as the third member of the MBIS team in 2017, and subsequently became an 80% HSLS faculty member / 20% GAC analyst in 2018 due to workload demand at the library.

According to the MBIS survey, 100% of respondants who had an in-person MBIS consultation are very satisfied (n = 40) or satisfied (n = 18). In-person consultations are preferred (63%, n = 74) over email (21%, n = 25), online/WebEx (11%, n = 13), or phone (4%, n = 5) consultations. Regarding the speed of response for phone or email consultations, 36% (n = 42) are very satisfied, 21% (n = 24) are satisfied, with only 1 respondant expressing dissatisfaction.

### MBIS Web Portal

MBIS maintains an active website [30] which serves as the digital gateway for HSLS bioinformatics services. The portal displays links to workshop guides, class calendar, MBIS blog/newsletter, consultation request form, and more. To provide a thorough overview of the MBIS software collection, we created an infographic embedded with links to guides dedicated to each of the listed tools [41]. These pages provide registration and access information, highlight key software features, and link to supportive webinars, tutorials, user manuals, and other relevant materials.

MBIS and the HSLS digital services team continually collaborate on HSLS-developed online resources such as data mining web tools and domain-specific search engines [42]. For example, a search tool prominently displayed on the MBIS webpage helps researchers via a federated seach to identify useful molecular databases/software, experimental protocols, lecture videos, and peer-recommended articles. The search engine, powered by IBM Watson Explorer[43] and licensed by HSLS, clusters the search results into meaningful categories for easy navigation.

In 2015, MBIS released “InfoBoosters,” an innovative, easy-to-install web browser widget that integrates digital text with databases to retrieve relevant information on demand [44] [45]. As life sciences research becomes increasingly interdisciplinary, scientific papers read by the research community often include genes, proteins, methodologies, and biological concepts outside of a readers’ domain of expertise. In order to thoroughly comprehend such articles, it is necessary to explore any unknown terms. Information is readily available in various molecular databases, but the reading format of journal articles (PDF or Web-based) does not provide links capable of accessing these databases directly from the article. The typical multi-step method to learn more is to (1) leave the article, (2) go to a separate online database, (3) search for the term of interest, (4) identify a reputable knowledge source, (5) scan it for the pertinent information, and then (6) return to the original article to continue reading. InfoBoosters improve upon this inefficient process and directly connect readers to databases such as *UniProt* and *NCBI* resources, as well as general information sites such as *Wikipedia* and *Vocabulary.com*. As an example, by highlighting a gene term in an article and then clicking on the “Protein-Info” InfoBooster, a pop-up window appears displaying protein-centric information for that gene as retrieved from the *UniProt* database. Infoboosters enhance user reading experience by immediately providing information at the point of need.

MBIS developed another search engine in 2016, called *search.bioPreprint* [46]. This tool helps researchers to comprehensively search preprint databases to discover cutting edge, yet-to-be-published or reviewed biomedical research articles [47]. Until the creation of *search.bioPreprint*, there was no simple way to identify biomedical research prior to journal publication, as preprints were not typically indexed in major databases, and were only discoverable by directly searching a variety of preprint server Web sites. This HSLS in-house developed tool offered a quick means of finding these types of articles and made an important contribution to the open-access open-source preprint movement.

MBIS survey results indicate that our web-based resources are used, although there is room for improvement. Two-thirds of respondants use some part of the MBIS website daily (7%, n = 8), weekly (29%, n = 34), or monthly (30%, n = 36), whereas one-third rarely (28%, n = 33) or never (7%, n = 8) use it. The vast majority indicate that it is very easy (16%, n = 19) or easy (69%, n = 82) to find what they need on the website, although a few indicate that is either not easy (8%, n = 10) or they have never used it (7%, n = 8).

### Outcome

With the growth of genome sequencing, the use of computational tools in biomedical data analysis is an integral part of the research [35]. However, due to limited time and training opportunities, many in the biomedical experimental research workforce lack the necessary skillset [10]. Thus, some researchers must either collaborate with expert bioinformaticians or outsource data analysis efforts to institutional bioinformatics core facilities. Given the limited availability of skilled bioinformatics professionals, this approach often requires extra time and expense for project completion. To speed up the process, researchers may prefer to perform their own bioinformatics analyses, and consequently require appropriate training.

MBIS fulfills such a bioinformatics training need. Our services are valuable to the Pitt research community thanks to a service-oriented philosophy as a library-based program available to all. The workshop format of 2-3 hours is particularly appealing over a more traditional semester-long for-credit bioinformatics class in that it (1) requires a lesser time commitment and (2) provides no-cost training to all members of a research team, including postdoctoral researchers and lab technicians.

Based on the survey results, the 17-year-old MBIS program is highly successful in providing bioinformatics software access, support, and training to the university research community. MBIS has considerably expanded in terms of software licensing, usage, and staffing since our establishment in 2002. From 2007 to 2018, the number of registrants for HSLS-licensed software has grown from 323 to 5435, the number of licensed software products has increased from 5 to 17, and the number of consultations has increased from 64 to 363. These heavily-used MBIS services are a reflection of the demand from the Pitt biomedical research community for sustainable bioinformatics support.

### Challenges

The maintenance of a library-based bioinformatics service is not without its obstacles. The primary challenge is keeping up with advancements in the field of bioinformatics; new technologies frequently replace older experimental methodologies, resulting in the perpetual emergence of new data analysis tools. Another considerable challenge is the management of the HSLS-purchased software licenses that have limited concurrent usage access. Users are occasionally denied access to software when the concurrency limit is reached, which is frustrating and results in an increased number of emails to MBIS staff with requests to troubleshoot. It is also demanding to create workshop content appropriate for all audiences, especially attendees with a diverse level of expertise and background knowledge. Condensing a convoluted, multi-step data analysis process, such as RNA-Seq analysis, into a three hour session takes a lot of time and care.

## Conclusion

Establishing and maintaining a library-based bioinformatics program with scalable infrastructure and sustainable service requires strong support from research leadership, along with dedicated funding for resources, infrastructure, and personnel. We believe that the benefits far outweigh the costs of running MBIS. The advantages to the library and the university are many: (1) significant financial savings via institution-wide network licensing of commercial bioinformatics software over individual researcher-based licenses; the negotiated institute-wide network licensing cost paid by the library is far less than the total cost if every software user paid market price [35], (2) continuous engagement with the research community through training and consultations increases library visibility and prestige, and (3) immediate access to quality software and associated training speeds up research and can prevent months of unnecessary, or flawed, analytical work.

We anticipate the demand for bioinformatics software and training offered through MBIS will continuously increase in response to easy access to the latest NGS sequencing and informatics technology provided by the recently established UPMC Genome Center [48]. Additionally, the recent shift of research interest from the characterization of heterogeneous cell populations to the high-resolution study of single cells will also multiply the need for large-scale data analysis [49]. As it has for the past 17 years, the MBIS will continue to adapt its educational outreach, collaborative partnerships, and licensed tool resources as needed to best support the Pitt biomedical research community.

We hope that our experience with the HSLS Molecular Biology Information Service may serve as a guide to other institutions and libraries interested in developing a similar program or expanding upon their current services.

## Acknowledgments

We thank Dr. Michael J. Becich, Dr. Uma Chandran, Dr. Fangping Mu, Nancy Tannery, Jeffrey Husted, Fran Yarger, and the members of the HSLS Digital Library Services team for providing assistance and valuable guidance in running MBIS. We are grateful to the late Dr. Michael Barmada for early support of our Next Gen Sequence analysis services in terms of software procurement, workshop instruction, and overall enthusiasm.

